# Systematic Nature Positive Markets

**DOI:** 10.1101/2023.02.13.528257

**Authors:** Alex Bush, Katherine Simpson, Nick Hanley

## Abstract

Despite decades of global commitments, and increasingly urgent warning of environmental instability, the demand for land to support economic production is still increasing. Isolated and disorganized actions will not be enough to avert ecosystem failures. As many developers are already required to compensate for their ecological impacts through restoration, many see markets trading biodiversity credits as a financial mechanism to counteract degradation and drive investment in conservation. The challenge stems from a desire to recognize the multidimensional nature of biodiversity that contributes to ecosystem integrity without making suitable offsets intractable to supply. Instead, most regulators have opted to streamline ecological assessment, and undermine ecological rigour, in favour of promoting offset supply and economic efficiency. As a result, all evidence suggests offset trading programs have so far failed to mitigate losses, let alone support “nature positive” outcomes. To overcome this disconnect, and support more effective and equitable biodiversity markets, we propose credits be defined by the *irreplaceability* of a site, a metric long-established in the domain of systematic conservation planning. Irreplaceability avoids the limitations of like-for-like trading, reduces costs of offsetting to developers and society, ensures farmers willing to sell are fairly rewarded for loss of earnings, and that sites critical to achieving conservation goals are safeguarded. We developed an ecological-economic model of a biodiversity offset market to demonstrate irreplaceability guarantees no net loss of biodiversity and is the most efficient metric for guiding investment toward the recovery of Nature.

## Introduction

More than 75% of the Earth’s land is degraded, and this has led to widespread biodiversity loss, undermining the well-being of billions of people, as well as our efforts to combat climate change (1). Current evidence suggests multiple planetary boundaries have been exceeded (2) and business-as-usual is highly likely to result in catastrophic collapse across many ecosystems (3). Despite this, numerous global commitments to reduce, stop or even reverse current rates of biodiversity loss have not been met (4, 5). Instead, reversing global terrestrial biodiversity trends will only be achievable if we adopt strategic, coordinated, and above all ambitious, action (6). To “bend the curve” toward a more nature-positive future, private sector funding of biodiversity conservation needs to be increased to complement longer-established publicly funded programs. The ENACT initiative (Enhancing Nature-based Solutions for an Accelerated Climate Transformation) launched at COP27 calls for the mobilisation of private finance to support action on nature and climate-related targets the world over, accompanied by robust environmental and social safeguards (7).

Nature markets are one tool to mobilise private finance to incentivise landholders to undertake more sustainable land management actions to enhance biodiversity (8). These markets create income streams (for example, in the form of tradeable credits) for landholders to undertake actions to protect and enhance a specified measure of biodiversity. Demand for these credits can be voluntary, for instance from individuals or companies who wish to offset their negative environmental impacts; or else are created through government regulation (9, 10). We refer to these regulated markets as “biodiversity offset markets”, where developers purchase credits to mitigate impacts on biodiversity due to impacts of development. Credits are supplied by landowners who switch their current land management (such as arable farming) to a more conservation-orientated alternative (such as wetland creation). By establishing an appropriate rate of exchange between sellers (landowners) and buyers (developers), biodiversity offset markets can, in theory, achieve no net loss of biodiversity or a net gain within some defined area at the lowest overall economic cost to society.

Within regulated nature markets, the choice of biodiversity metric plays a pivotal role in their ecological and economic performance (11). This metric establishes the units of trade, determining how a regulator or offset bank measures the gains in biodiversity resulting from restoration actions undertaken by landowners, and balances those against the expected biodiversity lost due to development impacts. Simple metrics based on a combination of the area and condition of habitat are often preferred by regulators (12, 13), easing the task of identifying matching biodiversity units, and assuming that habitat classes indirectly capture benefits on other aspects of the ecosystem (14). However, numerous studies have demonstrated that these approaches rarely benefit biodiversity in the manner intended (13, 15, 16).

In this study, we develop and then apply a new biodiversity metric for nature markets, derived from the Systematic Conservation Planning (SCP) literature (17, 18). SCP tools are designed to minimize the cost of achieving conservation targets. The importance of any specific site to achieving conservation targets is measured by its *irreplaceability*. A site that is essential to achieving targets is irreplaceable (and its loss could not be offset), whereas irreplaceability is low for sites which can be substituted for many others. Crucially, irreplaceability can aggregate the importance of a site for many biodiversity features, integrating the likelihood actions are successful across space, and ensuring overarching targets for the whole landscape are achieved. This integration represents a step change away from existing like-for-like compensation regimes in biodiversity offset markets (for example 19, 20, 21). Furthermore, if conservation targets are chosen to exceed their existing availability in a landscape, inherently these can only be achieved through restoration. This embeds net-gain as an implicit outcome, but only where this is needed to meet specific targets. We demonstrate that an offset market led by metrics derived from irreplaceability ensures the opportunity to achieve conservation targets is always protected, and the network selected is more economically efficient than using simpler offset metrics.

## Results

To answer whether irreplaceability offers us these advantages, we developed an ecological-economic model of a biodiversity offset market that ensures no net loss of biodiversity for the buying and selling of offset credits for a simulated landscape, using irreplaceability as the biodiversity metric and trading unit for credits. We assume a mechanism such as an offset bank or regulator which collects supply offers from all potential offset suppliers (farmers), in terms of their minimum willingness to accept (WTA) compensation for the offer of a specific offset credit. This minimum WTA depends on the opportunity cost of creating an offset in any specific location. We further assumed that the same mechanism collects demand offers from all potential buyers (developers), in terms of their maximum willingness to pay (WTP) for each offset credit. These maximum WTP values depend on the value of land in any location for development.

### Irreplaceability achieves conservation targets in an economically efficient manner

Our simulations demonstrate that using irreplaceability (α_sum_) guarantees progression toward conservation targets within an offset market (Fig. 1a). So long as potential gains from trade between buyers and sellers remain (WTP>WTA), all species eventually achieve their conservation targets. Economic gains from trade are initially high when trading first takes place (WTP>>WTA; Fig. 1b, note the log scale) but rapidly decline as more expensive and less irreplaceable offsets are required to meet demand. At every stage the market favours the greatest gains towards targets at minimal cost, favouring an economically-efficient solution. Conversely the irreplaceability metric strongly dis-incentivizes developments from taking place on land with high α_sum_ scores because the number of offset sites typically required to replace their loss is typically prohibitive (Fig. 1c).

**Figure 1.**
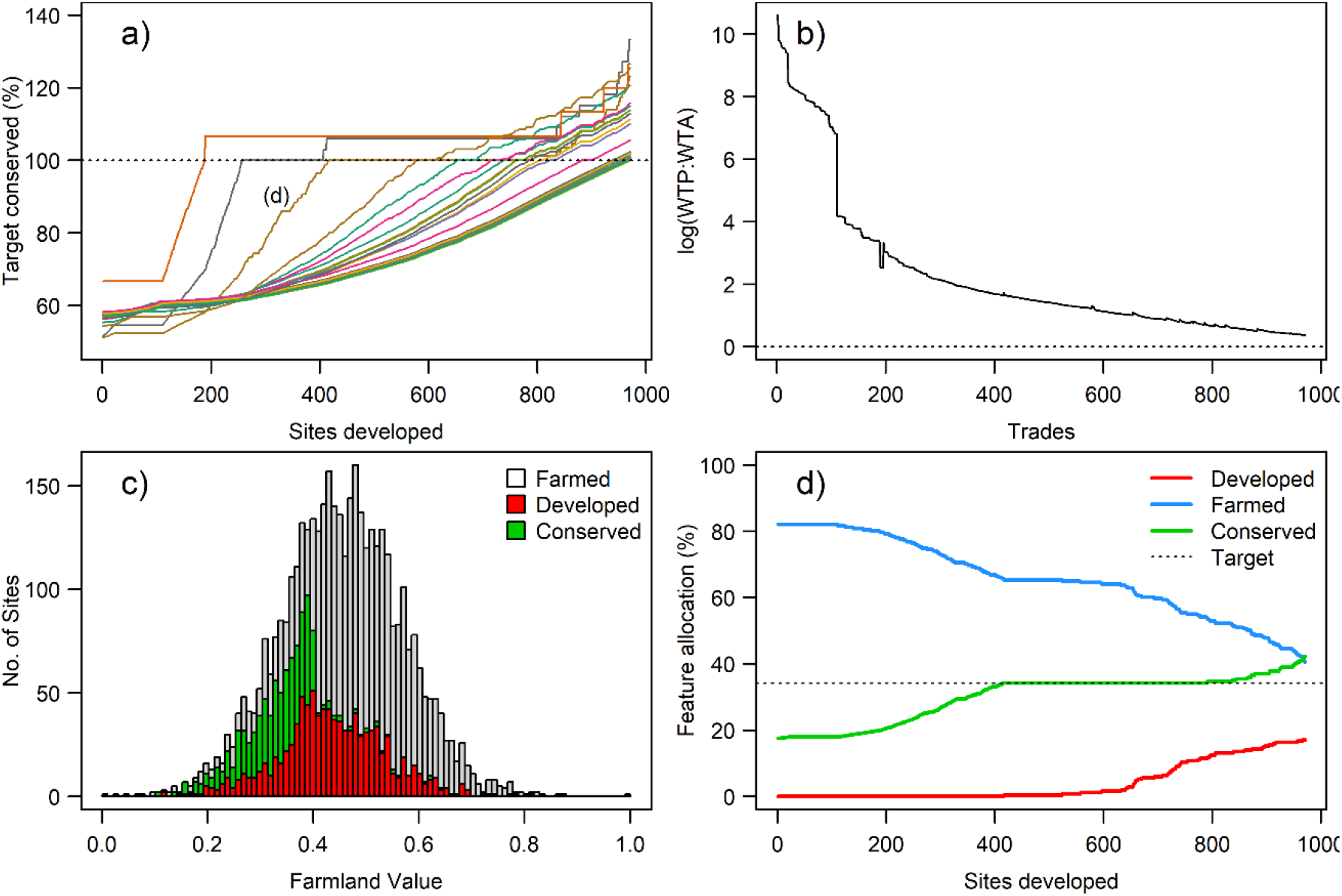
Example of α_sum_ irreplaceability offset market for 25 simulated species. Panel a) indicates the progress of each species toward its conservation targets (dotted line) as new developments requiring offsets take place. Panel b) illustrates the decline in the log ratio between Willingness-To-Pay (WTP) and Willingness-To-Accept (WTA) as representative of gains from trade, and c) displays the distribution of values for purchasing farmland in this simulated landscape, and the final proportion of those that were selected for development and conservation offsets. Panel d) displays the changes in the allocation of a single species (also identified in panel a) among land types as trading progresses.

Irreplaceability does not specifically prioritise sites that contain species rarely found in the landscape; it values sites based on the difficulty of achieving conservation targets without them. Nonetheless, as there are typically fewer opportunities to conserve rare species (i.e. low replaceability), sites that contain those species tend to score highly. Once a species target is reached (green line Fig. 1d), their contribution to the α_sum_ of remaining farmland is zero, meaning there is no benefit to its presence within new offsets, or cost associated with its occurrence at new development sites. Nonetheless, some species may eventually exceed their targets because they were present at offset sites added later to achieve targets of other species (Fig. 1a and d). As the α_sum_ contribution of species that have met their targets is zero, this reduces the burden for developers and increases their WTP at sites that contain species whose targets have been achieved (red line Fig. 1d).

### Accounting for more species in the market does not necessarily increase costs, or require more offsets, or a greater area to meet targets

The distribution of biodiversity, in particular the degree to which multiple targets are nested within the distribution of others, determines the degree to which additional sites are required to protect additional species. As illustrated by our simulations (Fig. 2), the network is specific to the assemblage, and how the ecological community correlates spatially with economic land values. Accounting for conservation targets of more species does not in itself increase the cost of conservation solutions, or require more trades, or more space to meet targets (Fi. 2b and 2c). However, in all cases the wide variation in outcomes for small subsets of taxa illustrates the risks associated with conservation policies reliant on small numbers of indicator species whose suitability to represent the conservation needs of biodiversity and ecosystem processes is unknown (22).

**Figure 2.**
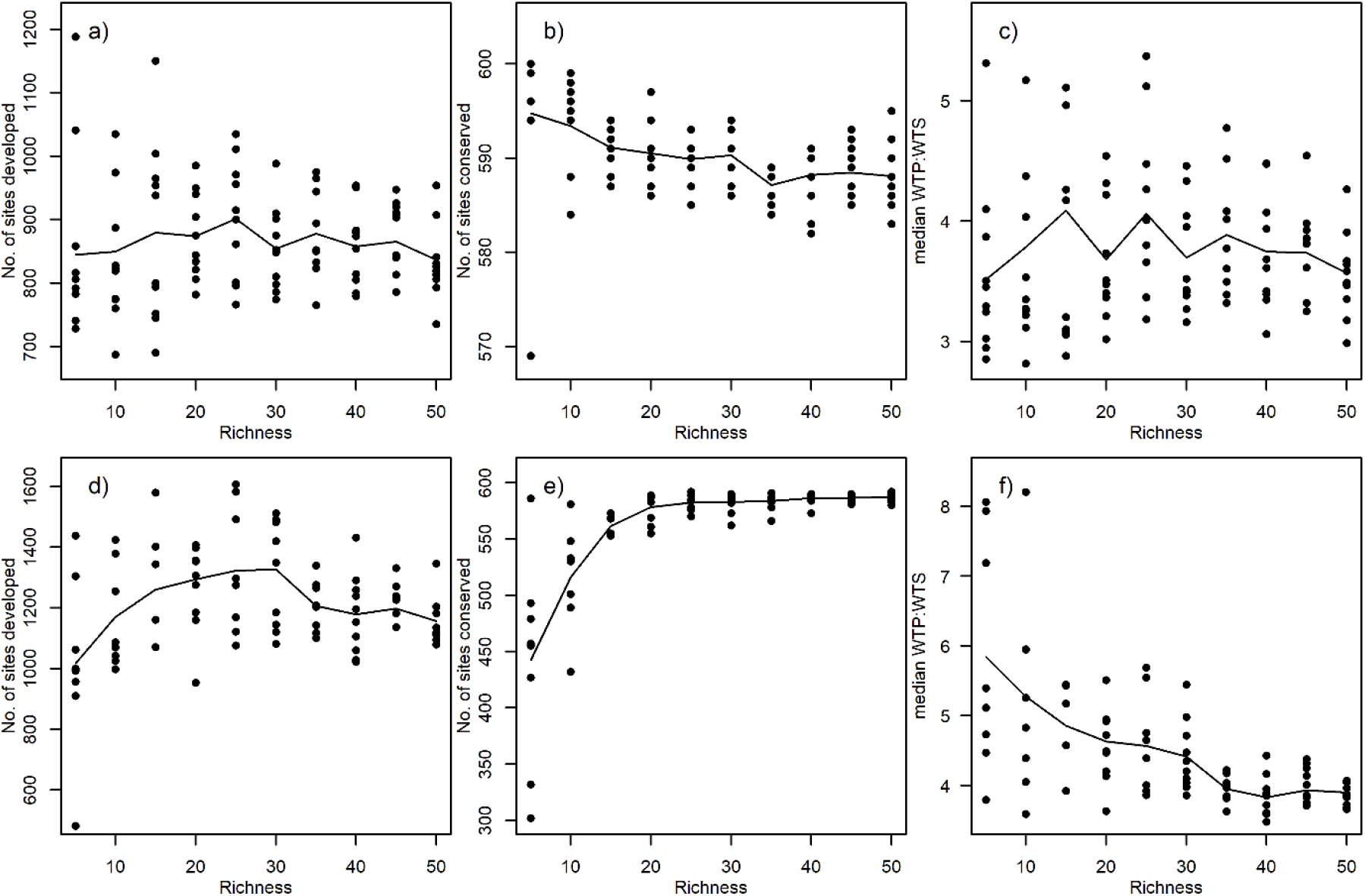
Variation α_sum_ irreplaceability market trading outcomes when the richness of communities is increased. Assemblages were drawn from communities of 200, with either a strong richness gradient (a,b,c) or no richness gradient (d,e,f). The columns show the number of sites selected for development (a & d), the number of sites required for conservation (b & e) and the median ratio between Willingness-To-Pay (WTP) and Willingness-To-Accept (WTA) (c & e). All conservation targets were achieved in each market simulation and lines of best fit were added based on local polynomial regression.

### Irreplaceability-led offsetting is comparable to optimal prioritisation

Site prioritisations generated by SCP are mathematically optimal, but rather than being reliant on land owners WTA, they assume that regulators or planners have full control over site selection and management. This is rarely the case where much land Is privately owned. If we assume developer’s WTP is sufficient to support continued trading, the α_sum_ irreplaceability trading market, like SCP, ensures all conservation targets are met; and as indicated in Figure 1d, it minimizes the cost to society given the constraints present at the time of trading. Indeed in our simulations the total cost of sites selected for conservation in an irreplaceability offset market was only 2-11% greater than using SCP, and this gap narrows as the numbers of species increases (Fig.3c) because the flexibility by which all targets can be achieved is reduced. Market solutions are more expensive when sites selected by SCP with low property values also provide high returns to developers, and therefore need to be replaced by an alternative complement of sites. The basis of SCP is that priorities are not simply the cheapest sites, or even the most ecologically diverse, but which site best complements and add to what’s already conserved. As a result, targets can be achieved by the market using very different networks of sites than the SCP solution (Fig.3b), and interestingly, those solutions may even require fewer sites in total (Fig.3a).

**Figure 3.**
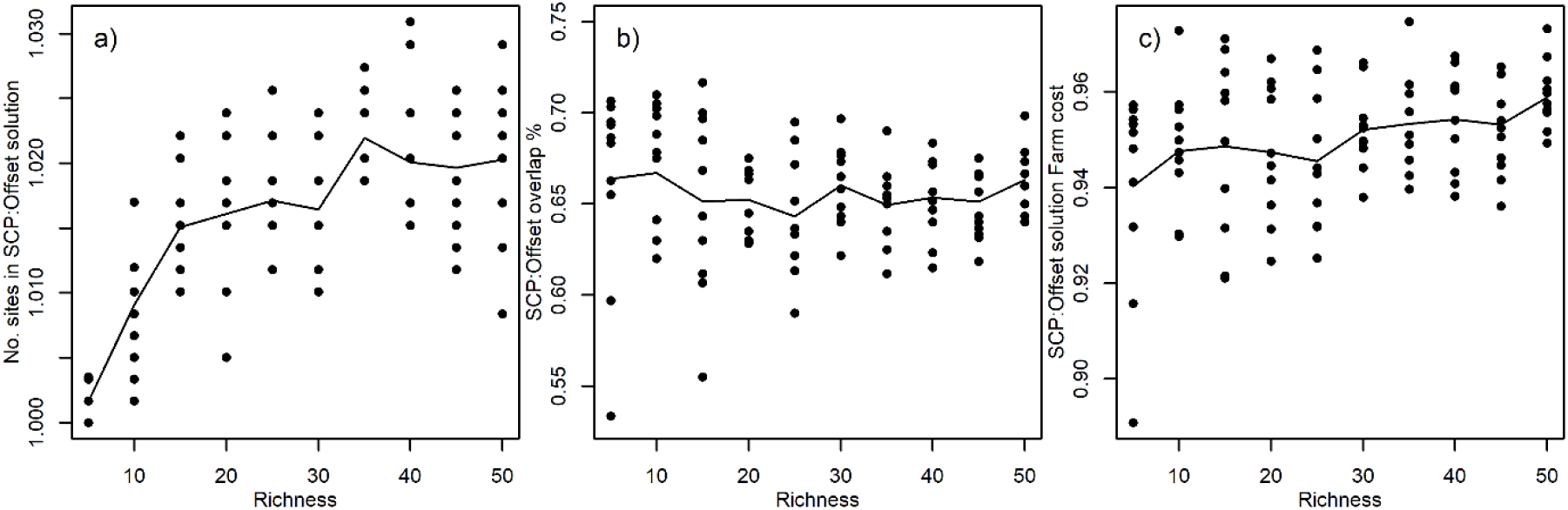
Illustration of a comparison between conservation networks selected by αsum irreplaceability market trading and “optimal” planning outcomes for simulated community with a strong richness gradient. Panel a) plots the ratio of network size when the richness of communities is increased; panel b) the percentage of planning units that are shared with the optimal network, and panel c) the ratio of network cost.

### Irreplaceability is ecologically and economically superior to simpler offset metrics

Markets where trade is governed by alternative simpler offset metrics (OM) typically failed to achieve all their targets (2%, 22% and 1% for OM1-OM3 respectively), even when property values were increased to support continued trading (Fig. 4). Offset Metric 2, in which sites are weighted by species rarity, was only more successful because targets in all our scenarios were directly proportional to their availability, and hence this was the only situation where fixed weighting could sometimes be appropriate. Yet the few occasions when alternative metrics did achieve all targets relied upon the subset of species selected to have narrow distributions which restricted the flexibility of selection. Where successful, solutions were achieved with a higher number of sites and at greater cost (115-130%), and none were successful for a larger number of species.

**Figure 4.**
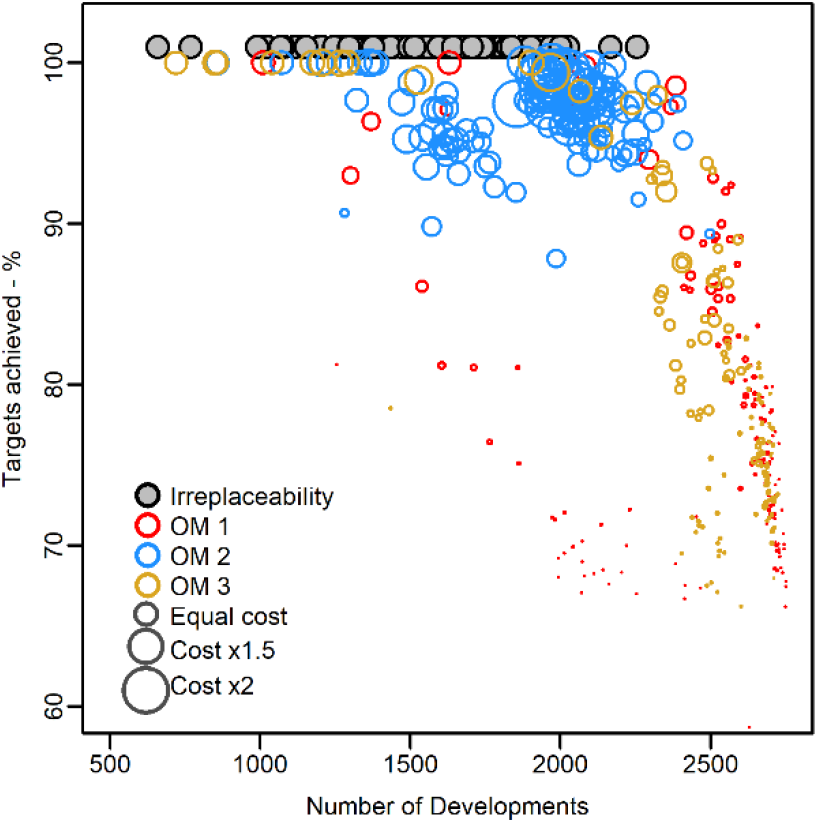
Comparison of conservation solution efficiency when guided by systematic conservation planning (SCP), or an offset market based on irreplaceability, and three alternative offset metrics described in the main text (O1-O3). Panel a) displays the total cost of farmland with the increasing richness of simulation scenarios, and panel b) displays the number of planning units that were Developed or entered into Conservation offsets. To make outcomes comparable only solutions that achieved >99% of targets are displayed. Note the solutions proposed by SCP are not associated with Development but are added to the plot to indicate the number of planning units conserved.

## Discussion

Land use is central to addressing global biodiversity conservation, as well as food security, poverty alleviation and climate change mitigation (23). The failure to coordinate appropriate and effective actions across sectors not only undermines commitments to drive a recovery of Nature, it further risks the sustained wellbeing of people. In this study, we have demonstrated that if relevant parties engage in trading of biodiversity credits based on a metric derived from irreplaceability, an offset market can support the most efficient trajectory towards conservation targets. That is, designing an offset market with irreplaceability as its metric delivers a low-cost way of meeting biodiversity targets.

Our approach challenges the current school of thought that to ensure no net loss (or achieve a net gain in biodiversity), “like-for-like” trading should be mandatory within a policy design (24, 25). Irreplaceability as a metric relaxes the need for equivalent species in every transaction and instead motivates restoration of species and ecosystems in greatest need (relative to targets), and where that action is most cost efficient. This element of prioritization ensures offsetting conserves the most important sites and at-risk species first, irrespective of whether they face direct development pressure. Previous research has hypothesised that increasing the complexity of offset trading metrics, in a similar vein to irreplaceability, is likely to reduce the number of trades and hence the economic efficiency of the policy instrument (9). In contrast, we demonstrate that simpler metrics are unlikely to achieve their primary goal and guide effective progress toward conservation targets, and that the economic cost of solutions based on irreplaceability were not dependent on the number of conservation targets we considered. In line with previous research, we demonstrate that the location of offset sites and overall cost of conservation actions is dictated by the overlap among ecological targets, and with ecological and economic heterogeneity across the landscape (26–29). Finally, if conservation targets exceed species starting availability because they anticipate restoration potential then net gain, rather than no net loss, is achieved at the market-scale.

The adoption of conservation planning tools allows conservation objectives to be achieved efficiently, but rather than implying the establishment of new reserves on former farmland, the intention is to value effective off-reserve management (30). Systematic conservation planning algorithms may define “optimal” solutions to meet all conservation targets, but in practice these networks are hard to implement when land is privately owned and landowner decisions are based on the relative payoffs from alternative uses (18, 31). By introducing regulations requiring developers to offset the predicted impacts of development on biodiversity the biodiversity offset market generates a positive financial return for farmers to invest in conservation that does not exist prior to this market being created. This study demonstrates that irreplaceability is an effective market metric to allow farmers and developers to independently engage in trades, while ensuring an underlying strategic approach is taken to secure the targets deemed critical to biodiversity conservation. Nevertheless, an ongoing problem in the successful implementation of biodiversity offset markets, and environmental markets more broadly, is the lack of regulatory capacity to implement the programme with an emphasis on the follow up monitoring of newly created sites (15, 32, 33).

### How can we avoid previous mistakes? Effective asset management requires monitoring

The quality of our knowledge of biodiversity is critical to estimating the appropriate allocation of land for conservation, arguing for resources, and negotiating trade-offs. To identify where a target can be achieved most effectively, irreplaceability credits combine knowledge of how ecological assets are distributed throughout the market’s jurisdiction, not just within sites associated with offset trading, and that information should be updated routinely to reflect their changing stocks. This is a potential challenge given monitoring have been cited as a key constraint to global action for many years (34), as well as prior attempts to organize biodiversity markets (15, 16, 35). However, a principle underpinning irreplaceability markets is that losses to development should not be sanctioned if they cannot be replaced, and in this context the value of ecological monitoring data gains new meaning. If our understanding of an ecological feature, like species distribution, is poor, we should err on the side of caution and protect a higher number of sites to be confident we have reached a target (36). Without this prudent approach, land and ecological assets upon which society depends may be lost before we have the knowledge to react. If caution due to data paucity overestimates the area required to achieve targets, this increases the difficulty of achieving targets and consequently the financial costs of offsetting for developers. It would therefore be in the interests of both market regulators and developers to improve monitoring to minimise the uncertainty of site’s α_sum_ irreplaceability, balancing the cost of further monitoring against expected efficiency gains for the market (37, 38).

Monitoring will be fundamental to the success of any financial market intended to support the recovery of Nature. It is key the market should represent as many asset types as possible, even if their distribution is uncertain, to avoid unintentional losses of biodiversity being permitted because those features were absent from α_sum_ calculation (39). Rather than rating performance according to the resources or finance committed, irreplaceability should reward landowners able to deliver ecological outcomes at low cost (34). The cost of monitoring has traditionally been prohibitive, but modern tools such as acoustics, molecular methods, automated imaging and remote surveys from drone and satellites have dramatically increased our ability to monitor many ecological systems at scale (40, 41). It is beyond the scope of this paper to provide an overview of these methods, but the capacity to efficiently verify restoration outcomes is growing, particularly if sampling design can be strategically adapted to minimise uncertainties in irreplaceability (42).

The biodiversity market is created by a demand for credits enforced by a regulator. The guarantee that conservation targets will be safeguarded and eventually achieved cannot be made if developers participation in offset trading is voluntary. The market regulator receives updates from monitoring sources to maintain oversight of each asset’s progress toward targets at the market-scale, thereby determining local site irreplaceability scores and the credits required for trades (35). The regulator is also able to intervene in the economic efficiency of the market, for example by subsidizing restoration costs on farms to increase the market supply of irreplaceability credits. While we recognize defining site irreplaceability based on the potential recovery of a site is challenging (43), include forecasting of the timeframe and risks (44, 45), those uncertainties are not barriers to adoption, rather motivations for targeted research (37, 38). Public support and trust will be strengthened by the transparency with which individuals can understand how local, and potentially highly visible, losses are accompanied by secure landscape gains designed to benefit society and the economy (46).

### Beyond biodiversity offset markets

Even with introduction of planning regulation, to avert substantial biodiversity loss and degradation of ecosystem services, we must raise our ambitions to begin restoring ecosystems (6). The resources available for conservation action are woefully inadequate compared to the resources invested in activities that further degrade or destroy nature (47), and yet the expected benefits of conservation investment far outweigh the costs (48, 49). The evidence of an ecological crisis is so serious that any action or investment is seen as positive, but this lack of discrimination also weakens the motivation of individuals and companies to support more transformative change. Irreplaceability credits can be used to recognize and reward private investment because they provide a comparable metric of performance within a market, even if two sites or actions impact different ecological assets.

Within an irreplaceability-market, an investor could anticipate the relative costs of their actions and define the performance of their investments in restoration and conservation for biodiversity in “net” terms. Irreplaceability could therefore be key to allowing fair recognition of investors’ contributions, while building public trust that companies statements of environmental responsibility match their claims.

The debates associated with pathways to sustainability and a nature positive recovery are highly value laden, “wicked” problems (23, 50), but we cannot expect ecosystem recovery to emerge from a piecemeal approach. Space is finite, both globally and within nations, and reconciling demands and interactions of complex multisector systems requires strategic oversight to avoid scenarios of ecological, economic and societal collapse (2, 51). Ecologists can identify what targets are required as a *minimum* to sustain species, ecosystem or process, but targets must ultimately be defined collaboratively with economists, social scientists, health economists and politicians. Incentivizing outcomes using systematic planning will become increasingly important as the collective benefits of multiple land uses diverge (52, 53). Irreplaceability would enable authorities to identify the targets and features that will pose the greatest conflict, and thereby accelerate the speed with which we can support Nature’s recovery.

## Materials and Methods

### Irreplaceability: recast for biodiversity offsetting

Systematic Conservation Planning is a rigorous, repeatable, and structured approach to designing new protected areas that efficiently meet conservation objectives (17). At an analytical level, the task is a classic resource allocation problem that either maximises conservation outcomes within a given resource budget or minimises the cost of achieving specified conservation targets (54). This structure has led to the use of SCP in supporting conservation decisions across the globe (18). A key strength of SCP is that it can incorporate a wide variety of data types, including attributes of ecosystems at all levels of structural, taxonomic, and functional organization, as well as accounting for social, financial and political constraints and opportunities (31, 55). The value of a specific site is based on its *marginal* contribution to achieving the conservation targets by complementing what features are already secured. A key feature therefore of SCP is that, unlike ranking procedures, properties of reserve systems emerge from the combination of areas, either through the complementarity of their composition or by their connectivity in space. This suggests a strong potential advantage for using a metric derived from SCP within biodiversity offset markets, where a need exists to be able to compare ecological gains and losses across space between development sites (where biodiversity declines) and offset supply sites (where biodiversity is increased due to the action of the landowner). Moreover, a biodiversity offset metric needs to make sense in the context of an overall policy target of no net loss or net gain in a specific aggregate indicator of biodiversity. This combination of an aggregate target with the need to compare gains and losses across space suggests that a metric or currency derived from SCP could have important advantages over the kinds of metrics investigated so far in the literature (26).

Provided with data on feature values for all planning units, planning unit costs, and the desired targets for protection, systematic conservation planning tools identify which sets of sites deliver conservation targets most efficiently (54). For convenience, we refer to “features” and “planning units” as *species* and *sites* hereafter. Often targets can be achieved by many different combinations of sites because alternatives exist with similar, or at least complementary, values. The importance of any specific site to achieving conservation targets is measured by its *irreplaceability*. A site that is essential to achieving targets is irreplaceable (and its loss could not be offset), whereas irreplaceability is low for sites which can be substituted by many others. An exact calculation of irreplaceability rapidly becomes intractable as the number of combinations to test scales exponentially with the number of planning units (56), and alternatives to estimate irreplaceability have been proposed (57). Most recently, Baisero, Schuster and Plumptre (58) proposed a new metric for describing irreplaceability (α) that defines the extent to which a site *k* is essential for achieving the conservation of species *s* as:

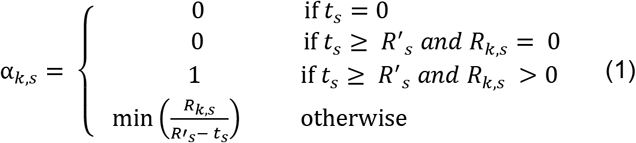

where the difference between the total availability of a species in the landscape *R’_s_* and its target *t_s_* indicates how much of that availability a site can contain *(R_k,s_*) before it becomes irreplaceable. Baisero, Schuster and Plumptre (58) define the combined site irreplaceability by taking the product of replacement probabilities (1-α_k,s_). However, this constrains site irreplaceability to between 0 and 1, and consequently no longer indicates whether a site was irreplaceable for one or many species. To retain this distinction and make comparisons among sites within an offset market equivalent, we use “summed α-irreplaceability”. We note Ferrier, Pressey and Barrett (57) also summed irreplaceability in their study for a similar reason, albeit with a different formulation for each species, and therefore this study specifically refers to the sum of α-irreplaceability (∑α_k,s_), which we abbreviate here to α_sum_.

### The Biodiversity Offset Market

The structure of the biodiversity offset market was based on the model developed by Simpson et al (2021). A single agent is controlling each land parcel, or site, who can decide to either develop their land for housing, generate biodiversity offset credits or remain in the current land use. For an agent to develop their land, each hectare acquired for new housing development requires a number of offset credits to be purchased from an offset provider equal to the measured biodiversity value of the site. The developer’s maximum willingness to pay (WTP) for an offset credit is determined by the expected value of land for housing development and the need to purchase offset credits. Ranking this WTP from highest to lowest yields a downward-sloping demand curve for offset credits. This WTP varies over space due to variations in house prices and the value of the site for biodiversity. We assume the offset credits are supplied by agents on agricultural land (“farmers”). Farmers change their current agricultural land management practices in a way which increases the biodiversity by a measured amount at the site. Every hectare given up to benefit biodiversity means one less hectare for agricultural production.

Furthermore, the farmer may incur restoration costs. Therefore, the conversion cost to the farmer consists of the opportunity costs of the foregone agricultural output plus any associated restoration costs. This is the farmer’s minimum price they will sell an offset credit for, known as their Willingness to Accept (WTA). Since agricultural productivity and profits vary across space (due, for example, to variations in soil quality or site altitude), the minimum WTA of farmers to create biodiversity credits will also vary. Ranking farmers from lowest WTA to highest WTA generate a supply curve for offsets. Farmers and developers interact in this market to generate an equilibrium, market-clearing price for offsets where marginal WTP and marginal WTA are equal, that is, where supply for credits equals demand for credits.

### Simulation

#### Inputs

To demonstrate the operation of a biodiversity offset market using the irreplaceability metric we simulated the probability of species occurrence within a 64 x 64 cell (or site) landscape. We used the R packages *NLMR* and *landscapetools* to control the degree of spatial autocorrelation in the baseline environmental gradient (59). Note however that α irreplaceability is determined by the global availability of that species, not their distribution, and that the simulation of maps was solely intended to communicate the parallels with field-data and empirical models. We subsequently simulated three communities, each with 200 species whose distributions were either equally distributed across the environmental gradient, or moderately and highly skewed towards one extreme to produce an overall gradient in richness (60). We ran offset market simulations based on subsets of species from each community, rising from 5 to 50 species and repeated 10 times each. More complex arrangements in response to multiple gradients are easily generated, but not considered further in this study. Likewise, in this study we did not account for lags or uncertainties in the restoration of offsets.

Four further pieces of information were generated for each site. The value of land for farming and for development were generated by defining their correlation to the environmental gradient (ranging from 0-1), although without a clear rationale for how these costs are expected to covary, both correlation coefficients were set to zero in our simulations. To reduce the likelihood that the market stalls when WTA<WTP which limits comparison among scenarios, development value was set to double that of farmland value. Next, each site is assigned to one of three initial land use classes: farming, conservation, and development in a 70:20:10 split. This represents the fact that landscapes already contain existing constraints, and the primary transfer that takes place in the market (i.e. as biodiversity is lost when farmed land becomes developed, so equivalent farmed land should enter conservation status). Lastly, a “habitat” layer is generated to indicate where habitat, and hence species, currently occur on farmland and conserved sites to define the baseline from which “gains” should be compared. Farmland sites without habitat, but suitable environmental conditions for species to occur are treated as areas with restoration potential. The final inputs are the conservation targets for each species. To illustrate a scenario of net-gain, rather than no-net-loss, we set targets in all scenarios to be the equivalent of each species existing availability, plus 20% of their restoration potential at farmland sites.

#### Market

At each stage the α_sum_ irreplaceability is calculated for all sites. A farmland site that is not irreplaceable for any species and has the greatest WTP (£/α_sum_) is selected for development. If the Development site α_sum_ is 0, either because the site has no species potential at all, or because all species with potential have already achieved their targets, then no offset is required. Otherwise, an offset site with the lowest WTA (£/α_sum_) is selected and either all, or a fraction of species values at that site are assigned to conservation status. The species values at the developed sites are removed from the global total *R’_s_* and the values added by the offset are deducted from the remaining targets *t_s_*. These steps are then repeated until all species conservation targets have been achieved, or there are no mutually beneficial opportunities to trade in biodiversity credits remaining (that is, a market equilibrium where for all remaining sites WTP<WTA).

#### Performance

To rate the performance of an offset market based on α_sum_ irreplaceability we compared the efficiency with which targets were achieved using alternative metrics for the same landscape. Firstly, the R package “prioritizr” was used to identify the exact optimal combination of sites that achieved all conservation objectives for minimal cost (61). Secondly, the offset market was re-run using three alternative site-based metrics that increasingly reduced the need for the information involved in strategic planning. The first offset metric (OM1) weighted site scores by the inverse of each species range, thereby favouring the rarest taxa in the landscape (62). OM1 scores were also continually updated to reflect changes in global availability due to the market. OM1 assumes the same degree of knowledge as required for the α_sum_ irreplaceability, but without setting targets. Updates to planning unit scores reflect species’ global availability, but not complementarity to areas already protected. Offset metric 2 is equivalent to OM1, but values for each planning unit are not updated over time meaning weights for each species were fixed at their starting value. This metric required the same initial understanding of species distributions but does not require a register of species affected by previous offset transactions. Finally, offset metric 3 (OM3) was based solely on how many species were present, but not as above, by which species, meaning only a map of species richness would be required to guide a market.

The code and a full description of the results reported in the paper are provided in the supplementary material.

## Acknowledgments

The authors were supported by UK Natural Environment Research Council funding (natfin10004).

