## Supplementary Material 1 for "Systematic Nature Positive Markets"

### Supplementary Material
### Systematic Nature Positive Markets: offsetting with irreplaceability

Bush A., Simpson, K. and Hanley, N.”

Dec 2022

**Offset Market Simulation** **6**

**Results of a single simulation** **7**

**Results of all simulations and scenarios** **11**

**Comparison with Systematic Conservation Planning** **12**

**Comparison with Non-Systematic Offset Metrics** **15**

### Introduction

Our study focuses on a hypothetical region in which the landscape can be divided into three mutually exclusive land-use types: farmland, built developments, and conserved habitats. The profit a developer can achieve is based on the value of a site after development minus the cost of purchasing that land. More recently legislation governing developer's responsibility to ensure No-Net-Loss (**NNL**) or Biodiversity Net Gain (**BNG**) to receive planning approval has become more common. NNL and BNG policies require developers to mitigate their environmental harm by contributing to conservation offsets proportionate to the impact the new development will have. Therefore for a site to be a commercially viable opportunity for a developer, the value of the development must not only exceed the cost of land, but also the cost of delivering biodiversity offsets. As a result a developers **Willingness-To-Pay** (WTP) to develop depends on how many offset units are required and the financial cost per offset unit. The financial cost of delivering offsets depends on the price at which farmers are **Willing-to-Accept** (WTA) offset credits, which they can create by switching from farming to conservation. This study proposes the currency of site value for biodiversity is best defined by a metric that integrates the priority of all conservation targets, thereby systematically promoting net gain and guarding against unintended consequences. For a more complete description of the metric, namely the sum of alpha-irreplaceability, please see the main text.

This document describes in greater detail the simulated landscape and offset market conditions we used to demonstrate the advantages of an offset metric based on irreplaceability. Users can simulate ecological communities and landscapes as we have below or provide the outputs of their own empirically derived ecological models to explore how land values might be expected to alter the outcomes of ecological offset markets in their region.

---

### Simulated Inputs

The following packages were used to simulate landscape and species as feature data:

```
rm(list = ls())
library(NLMR)
library(raster)
library(RColorBrewer)
library(virtualspecies)
```

#### Landscape

The first step in our simulation is the generation of a habitat suitability layer, scaled from from 0-1 (**Fig.S1**). Note however that currently the irreplaceability score i not spatially explicit because it does not account for connectivity among sites. Nonetheless, we chose to generate a landscape with spatially autocorrelated gradients rather than a table of data to draw a more direct comparison with the underlying geographic layers that are typically used to predict species occurrence. All simulations reported below used the same landscape that contained 4096 planning cells (64 x 64 cells).

```
Dim  = 64
N    = Dim^2
Env1 = nlm_fbm(Dim, Dim, fract_dim = 1, user_seed=2785)
```

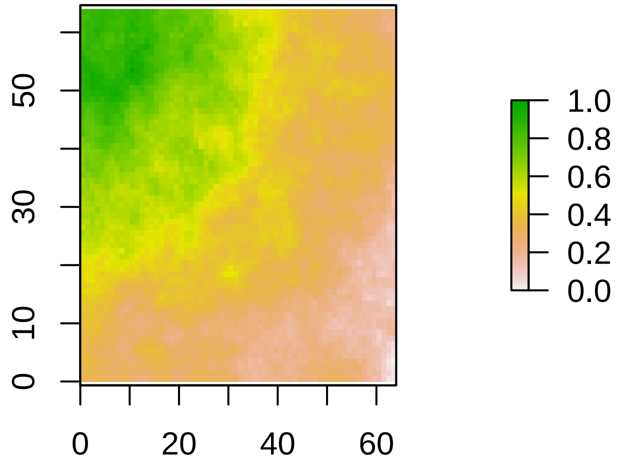

**Figure S1:** Simulated abiotic landscape.

#### Species

The **virtualspecies** package offers a multitude of ways to simulate species with more or less complex environmental responses. Nonetheless, for the purposes of this study we adopted the simplest approach available and described niches as normal response functions. To demonstrate the ability of systematic approaches to combine attributes of any number of conservation features we generated 500 virtual species to create communities. To create communities with increasingly strong gradients of richness and thus differences in conservation value, we drew values from three beta distributions to skew the value of species optima (i.e. the mean of normal response)(**Fig.S2**). Typically the distribution of species prevalence is skewed towards rare species with narrow ranges (specialists), and relatively few species are present across a wide range of environmental conditions (i.e. generalists). In a degraded landscape where much of the original natural habitat has been lost, remaining fragments becoming increasingly likely to represent an irreplaceable resource to one or more species. However, to illustrate how we should compare ecological value for less specialist species in an offset market and ensure the majority of land cells had at least the potential to provide habitat for some species, niche standard deviations were drawn from a beta distribution ($\alpha$=1, $\beta$=11) that allowed species to occur on average in 66% of cells.

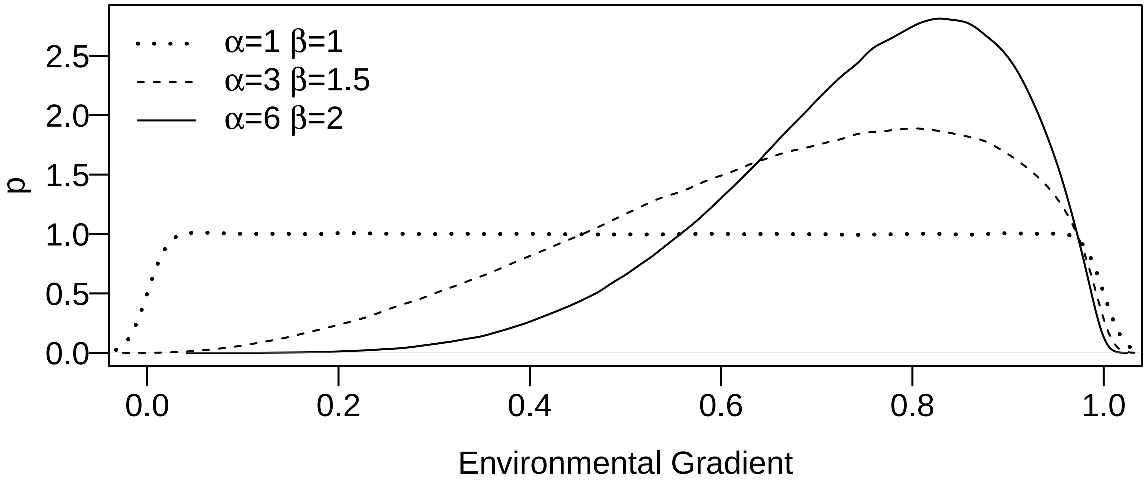

**Figure S2**: Distribution of species niche optima, and thus overall species richness, across the environmental gradient for three simulated communities.

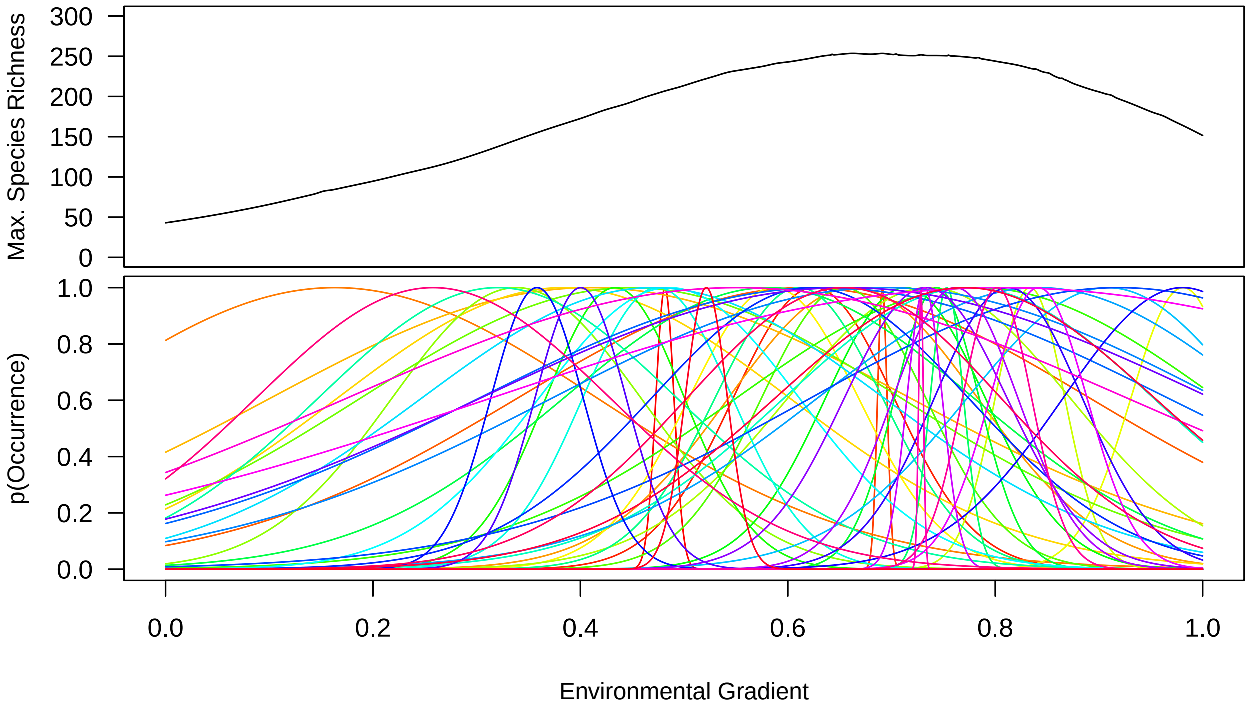

**Figure S3:** Fifty of the 500 species in a community simulated with a moderate gradient in species richness ($\alpha$=3, log($beta$)=1.5).

The scenarios result in communities that display contrasting distribution of species richness across planning units (**Fig.S4**), and within each community the distribution of species occupancy means that some will be rare and others common (**Fig.S5**). In general we expect representative conservation outcomes to be easier to achieve when priorities for one target overlap with those of others, which is potentially more likely when strong richness gradients exist or species have broad tolerance and occupy many sites.

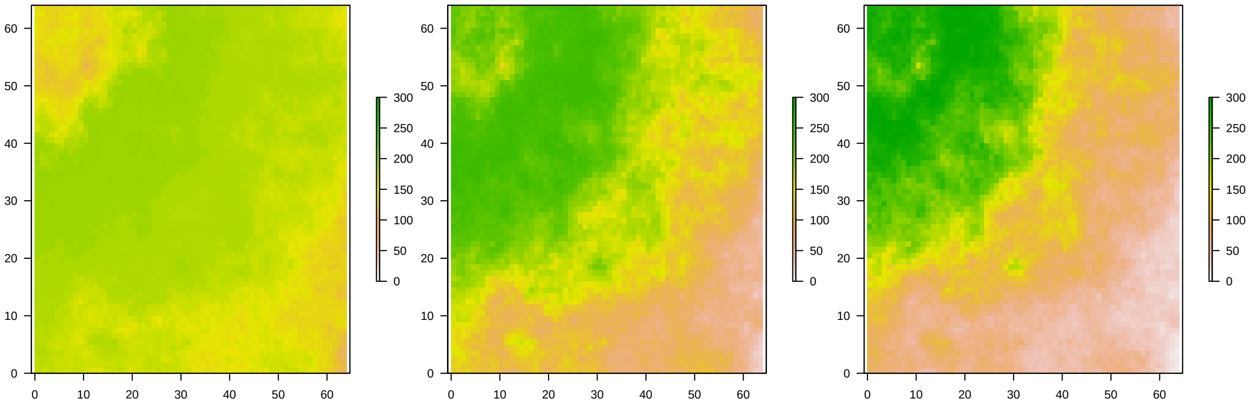

**Figure S4:** Distribution of species richness in simulated landscapes.

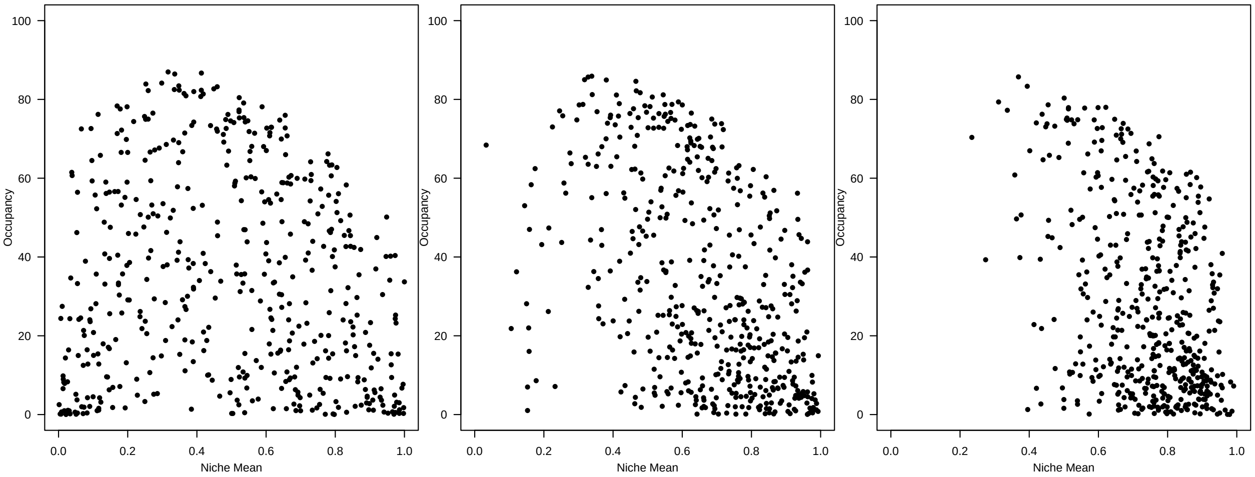

**Figure S5:** Distribution of species occupancy in simulated communities.

#### Land Use and Land Value

Farmland value (*Farm.value*) represents the agricultural income foregone by a farmer when their land is used for development or conservation offsets. It is thus a measure of opportunity cost to each farmer in the landscape of giving up current farm output and switching to an alternative land use (housing or conservation). To calculate the efficiency of biodiversity features per financial unit this value is constrained to be positive (strictly >0). The layer was defined according to the degree to which farming productivity is correlated with biodiversity potential:

```
# Load function to define layer correlation
source("define_correlation_of_new_vector_function.R")
# Create duplicate spatial layers
Dev.value.map = Farm.value.map = Env1

# Start with no correlation
Farm.value.map[1:N] = newx.defined.correlation(x1=t(Env1)[1:N], rho=0 )
Farm.value = Farm.value.map[1:N]

# Add a small amount to Farm.values so they are not zero
Farm.value[Farm.value<0.001] = 0.001
```

Development values (*Dev.value*) reflect the financial value of a site once it has been developed for housing. This is specified as varying spatially, and may correlate to some degree with farmland suitability for instance. The degree to which *Farm.value* and *Dev.value* co-vary with biodiversity values is important because they strongly influence the cost of supplying offsets capable of achieving conservation targets, and thus the willingness of developers to pay. Nonetheless, while we can manipulate the distribution of biodiversity value in the landscape above, and the correlation of *Farm.value* and *Dev.value* to those gradients, we do not have a firm understanding of how farmland and development values are expected to relate to one another. Therefore, for the time being *Dev.value* is assumed be to not be correlated with *Farm.value* (and therefore, in this case, also uncorrelated to the environmental gradient).

The definitions above for farm and development value greatly simplifies offset provision. The market is simulated on the basis of successive offset trades, but there is otherwise no element of time (our model is thus akin to a one-shot game). In this demonstration the simulation considers conservation offsets to be a permanent arrangement, and hence the farm income forgone by the farmer are the same as if they had sold their land to a developer. In reality we understand farmers may only wish to enter fixed-term arrangements to deliver offsets which may reduce the costs, but will in turn also moderate the expected value of biodiversity gains at an offset site, particularly in the context of permanent losses at a developed site. The lack of timescale also means our simulation considers the conversion from farmland to an offset to instantly guarantee secure conservation gains, whereas often the lag between restoration starting and biodiversity potential being achieved can be considerable. Importantly if there is an added cost to rehabilitate a site and maintain that restoration, these costs are not included in addition to property value in *Farm.value*. These assumptions are important, but do not alter way the market functions because this primarily depends on the ratio between site's *Farm.Value* and *Dev.value*. Therefore within the context of a simulation the consequences of different relative property values can be easily explored. As one of the primary interests of this study was to compare an optimal approach using systematic conservation planning with the outcome delivered by an offset market, we wished to see the market achieve all targets. To ensure the market continued we ensured demand for offsets was high by doubling *Dev.value* so that exceeded *Farm.value* in most cases.

```
Dev.value.map[1:N] = newx.defined.correlation(x1=t(Farm.value.map)[1:N], rho=0 )
Dev.value.map = Dev.value.map * 2
```

Finally, as we mentioned at the start, we consider the landscape to include three land-use types; farmland, built developments, and conserved habitats. The goal of the offset market is for a target amount of each conservation feature to be protected within the conservation land-use category, and this is therefore expected to build upon and complement the areas of the landscape already under conservation management. Likewise we include existing development simply to emphasize that the approach must work around existing constraints (for example, developed land cannot be converted into conserved land), and we can block those areas out from consideration. For the purposes of this demonstration we split the landscape into clusters based on a ratio of 70:10:20 for farmland, developed land, and conserved sites respectively (**Fig.S6**).

```
landuse = nlm_randomcluster(Dim,Dim, p=0.1, ai=c(0.2,0.7,0.1), rescale=TRUE)
```

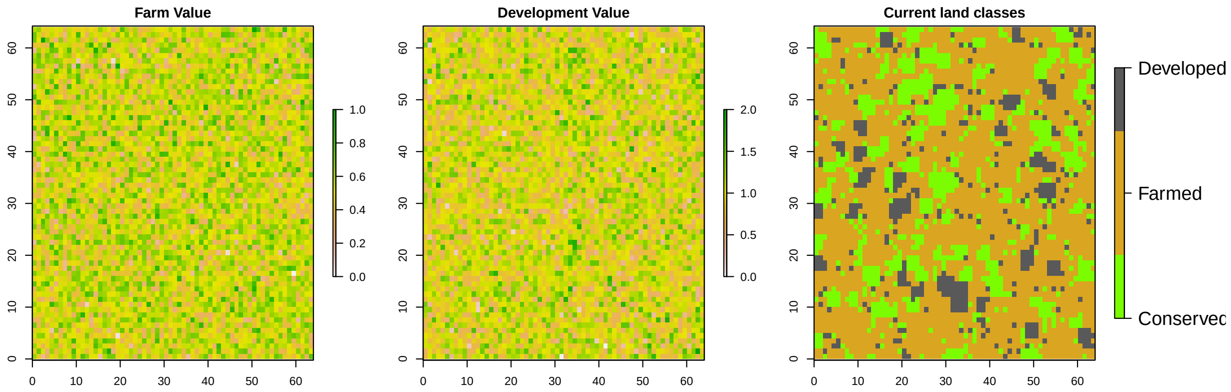

**Figure S6:** Maps illustrating the distribution of *Farm.value*, *Dev.value* and the starting type of the three land-use classes.

All simulated inputs are saved at this point so that subsequent market simulations only vary the identity and number of species under consideration.

---

### Offset Market Simulation

At this stage we wish to draw attention to the *proxirr* package released in conjunction with the study by *Baisero et al. 2021* which described the derivation of alpha- and beta-irreplaceability for conservation planning. The following code does not require installation of their package, but the function for alpha-irreplaceability was adapted from their work; the main modification relating to the summation of alpha-scores across taxa at each site to balance offset comparisons among sites that contain multiple irreplaceable, or near irreplaceable, features.

```
## Warning: package 'raster' was built under R version 4.1.3
```

```
## Warning: package 'sp' was built under R version 4.1.3
```

```
## [1] "citation('proxirr')"
```

```
## [1] "Baisero D., Schuster R. & Plumptre A.J. Redefining and Mapping Global Irreplaceability. Conserva
```

To load the code for simulating the offset market, source the function for **irreplaceable.market.sim**.

```
source("irreplaceable.market.sim.function.R")
```

In this study we include comparisons of the offset market equilibrium with '*optimal*' solutions generated by a systematic conservation planning (SCP) algorithm. Our function contains the functionality to run a *prioritizr*, but please note that *prioritizr* has its own dependencies, including the external optimization software *Gurobi* (Gurobi Optimization LLC, 2021, https://www.gurobi.com). If you wish to compare with the systematic conservation planning solutions, please refer to instructions for installing this package and how to access Gurobi.

```
library(prioritizr)
```

As well as comparison with a mathematically optimal outcome obtained using a systematic conservation planning algorithm, we also compared the performance of the irreplaceability-market with other increasingly simple metrics of biodiversity value at a site (as used, for example, in Simpson et al, 2022), and the equivalent function for simulating those outcomes is included in the code provided.

```
source(irreplaceable.market.sim.function.R)
```

We ran the market simulations 20 times each for random subsets of the 500 species at seven levels of increasing richness. Conservation targets were based on the simulated probabilities of species' occurrence and we chose to set targets in such a way to guarantee the delivery of net gain. The first part of the target is based on preserving the existing "stock" of a species in a region by selecting the values for that species in all existing conservation sites, essentially achieving no net loss. The second step to setting our targets added a fraction (20%) of the species potential in farmland, which is inherently additional to what currently exists and thus represents a net increase for all species. We also explored outcomes when targets were consistent or variable among species but, relative to the variation among simulations due to the identity of the subset of species selected, the balance of targets proved to have a minor effect. In practice targets would not be selected as function of potential; instead stakeholders would propose specific targets that better reflect ecological needs for species persistence as well as societal needs based on ecosystem functioning. Ideally these targets would also apply the precautionary principle so that if mistakes are made or sudden changes occur, irreversible losses due to species' extinctions or major habitat transitions do not occur before those features have the opportunity to recover. Finally it is worth noting that although we demonstrate irreplaceability is an efficient offset metric, higher targets come at greater cost. If we had been interested in the sites selection for even more ambitious targets, we would also have needed to consider increasing the ratio between development and farmland values in order to ensure supply to the market.

---

### Results of a single simulation

It is possible to plot the progress of all species towards their conservation targets during the simulation (**Fig.S7**). When all features have reached their targets the function ceases the plot clearly demonstrates how after each new development, and thus new offset added to the conservation land-use, the total stock of species moves closer to achieving their respective conservation targets. Although the irreplaceability score is site-based, those species for whom the targets are most difficult to achieve are effectively prioritized first because the lack of alternative sites to conserve them means that those sites where they are found receive higher irreplaceability scores. This means that farmers have a greater incentive to conserve these more valuable sites, since they will attract a bigger reward in the offset market. Once a species has reached its target it no longer contributes to irreplaceability scores, either in calculating development impact or offset delivery. However, their total may continue to rise (i.e. exceed 100%) because they are also present at other sites that are required to meet conservation targets of other species.

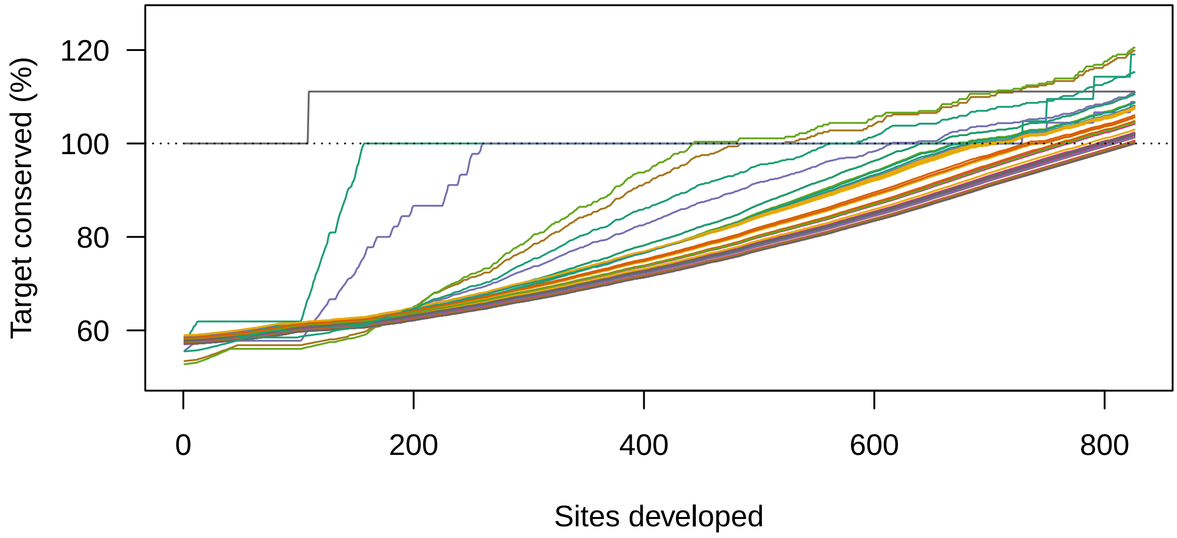

**Figure S7:** Progress of 50 species toward conservation targets followig successive offset trades. Market simulation ceases when all species have achieved their targets.

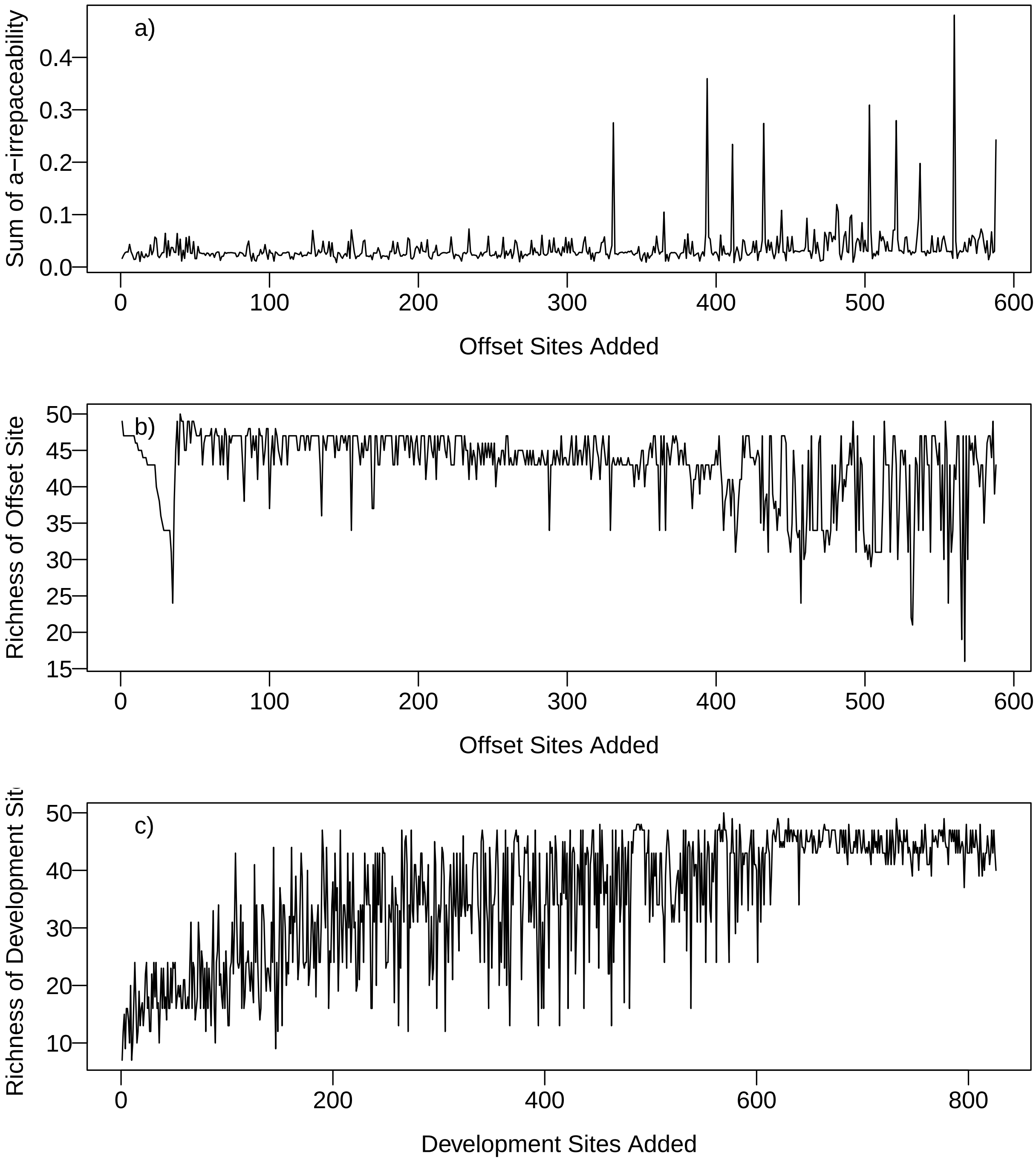

**Figure S8:** Plots illustrating the variation in sites selected by an offset market simulation based on 50 species. Panel a) displays the sum of alpha irreplaceability of selected offset sites, panel b) the richness of selected offset sites, and panel c) the richness of developed sites.

The sum of alpha-irreplaceability at any given site, its relative value to achieving conservation targets, is continually changing as developments and offsets remove features from consideration or remove the necessity to achieve gains elsewhere. As such it is difficult to interpret the irreplaceability value of offset sites (**Fig.S8a**), particularly as their selection interacts with farm and development property values. However, in the example provided there is a clear trend in the richness of offset (**Fig.S8b**) and development sites (**Fig.S8c**). Initially there is a sharp decline associated with the requirement to conserve the rarest species first, but after this the focus switches back to selection of the more diverse sites that provide broad benefits, and gradually declines over time. Conversely we expect the sites most attractive to developers will have least biodiversity value because this reduces the costs of delivering corresponding offsets, and indeed the richness of developed sites is initially very low and then rises. Further note that while developments may be forced to select increasingly diverse sites, in this simulation 14 species had met their targets after about the 700th offset trade (**Fig. S7**), and hence there would no longer be any penalty for development that impacted sites where those species were present. Another example of this is evident for those species that meet their targets relatively early in **Fig.S9**. At first, potential within farmed sites is converted to gains within conservation, but once their targets have been achieved, and there is no longer a penalty for their loss, it is evident that many more of those farmland sites begin to be selected for development.

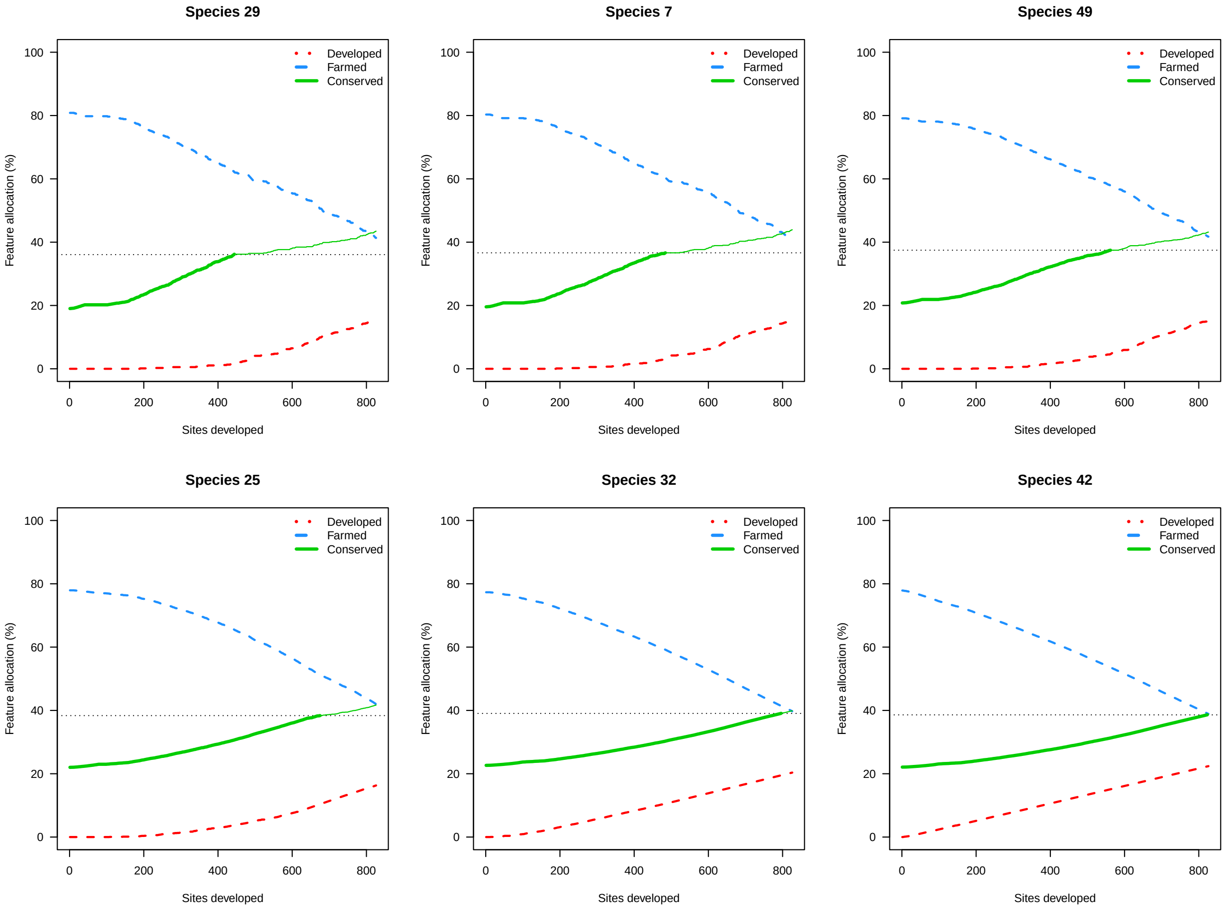

**Figure S9:** Plot of 6 species in the order they achieved their targets illustrating how the total habitat potential for each species was reallocated from farmed to developed or conservation land-use classes as the offset-market progressed. The dotted horizontal line indicates the target for each species and once this has been achieved the green line is thinner because the potential for that species is no longer considered in development impacts.

Finally, as well as plotting how ecological values change during the course of the market, we can also track changes in the ratio between developers' Willingness-To-Pay and farmers Willingness-To-Accept. The rapid decline in the ratio indicates that relatively few locations in the landscape had a high margin between their development value and the farmland value that subsequently increases the WTP. Nonetheless, in this particular case there were a large number of sites where this ratio was still sufficiently high that trading continued until all targets were achieved (note that, so long as the maximum WTP of at least one developer exceeds the minimum WTA of at least one offset supplier, then gains from trade exist, so the market will not yet have reached equilibrium). Our offset market "clears" (reaches equilibrium) when all these potential gains from trade are realized - when supply equals demand.

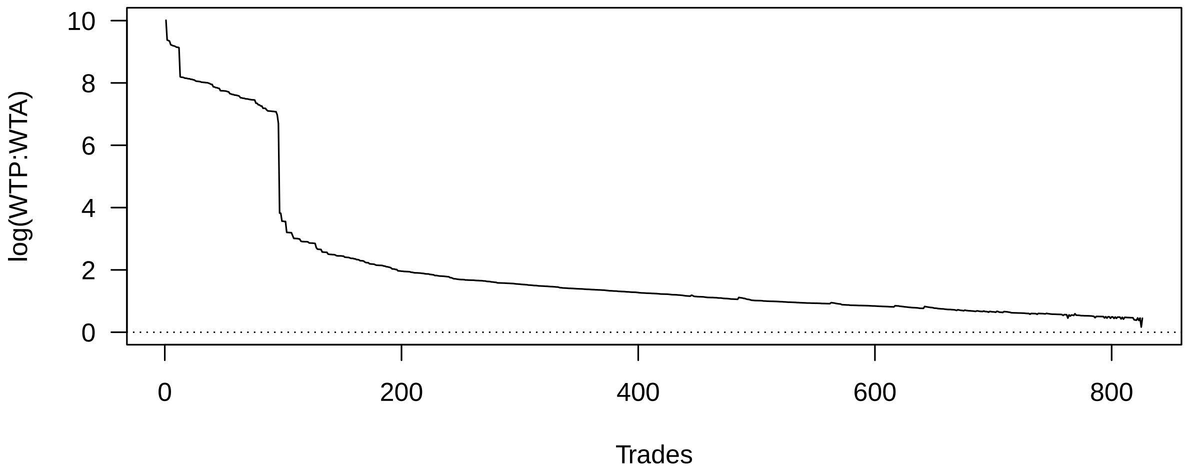

**Figure S10:** Example of decline in the log ratio between Willingness-To-Pay (WTP) and Willingness-To-Accept (WTA) during offset market simulation. If the ration between WTP:WTA had reached 1 (0 on log scale), there would no longer have been sufficient value to justify development and the market would have ceased.

---

#### Results of all simulations and scenarios

It can be instructive to simulate how sensitive markets are to the inclusion or exclusion of different features, or to vary the respective land values for farming or development. However, it should be no surprise to realize that the outcome of the offset market, including the number of developments, number of sites allocated to conservation, and the financial margins that ensure adequate supply are inherently tied to the scenario simulated, and hence if we alter the cohort of species considered or their distribution we get different patterns of results. As such the three landscapes scenarios we generated are simply illustrative and no firm guidelines or predictions can be made fm them. We would expect the variability in outcomes to decline as the number of species included in the market increased (left to right in each panel of **Fig.S11**) because fewer combinations of sites would be capable of achieving all targets. Conversely there was little difference in the overall cost and number of trades taken to complete markets in landscapes that had very different distributions of richness within them (comparing rows within **Fig.S11**), most likely because the subset of species selected didn't necessarily represent the underlying differences.

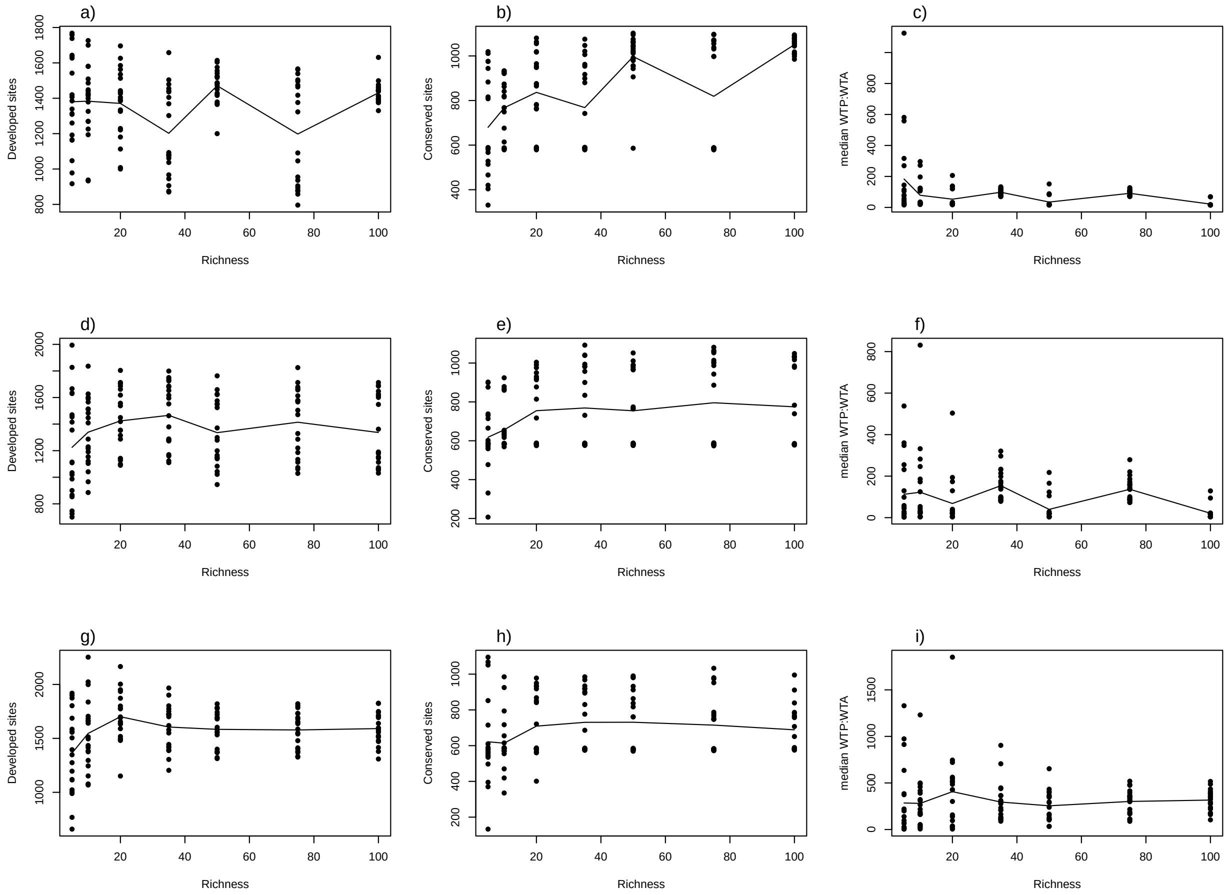

**Figure S11:** Results of offset market simulations for an increasingly large subset of species drawn from landscapes that featured an evenly distributed (a, b and c; $\alpha$=1, log($beta$)=1), moderately biased (d, e and f; $\alpha$=3, log($beta$)=1.5) and strongly biased (g, h and i; $\alpha$=6, log($beta$)=2) gradient in species richness. The first column (a,d,g) displays the number of developments that were required in each simulation before all conservation targets were achieved. The second column (b,e,h) displays the number of sites required to achieve targets for increasing numbers of species, and the third column (c,e,i) shows the changes in developers mean profit margin (ratio of developer's Willingness-to-Pay (WTP) to farmer's Willingness-to-Accept (WTA)).

---

### Comparison with Systematic Conservation Planning

Systematic Conservation Planning (SCP) identifies the optimal allocation of resources to achieve a goal (our conservation targets) given particular constraints (minimize the total cost of foregone farming income on land that would need to be set aside for conservation). An SCP solution therefore represents the best-case possible for an offset market because developers would only be paying the absolute minimum additional cost necessary to offset the impact their development has on achieving conservation targets. For an economist, the SCP solution is the *social planner's optimum land allocation.* **Fig.S12** shows that the manner in which conservation targets were achieved using offsets by the proposed irreplaceability market closely approximate those of the optimal SCP outcomes. Firstly, **Fig.S12a** shows that while the approaches selected a similar number of sites (i.e. ratio is near 1), for higher numbers of species the SCP solutions often chose a slightly higher number of sites to achieve all targets. While this result may appear counter-intuitive, **Fig.S12b** shows that even in this relatively small simulated landscape there was only an overlap in the sites dedicated to conservation for approximately 60% of cells (see also **Fig.S13**). While minimizing total area is often a good proxy for minimizing cost, **Fig.S12c** shows that the SCP solution was achieved on farmland that was 8-11% cheaper overall, even when it included more sites than the irreplaceability market. Conversely, **Fig.S12d** indicates that the SCP solution also selected land with higher potential development value than the offset market, effectively because this competing land-use is not considered as an additional constraint. We can therefore infer that in these simulations, where development and farmland cost were uncorrelated, some of the cheapest farmland on which to achieve conservation outcomes also carried significantly greater value to developers and hence during the allocation of offsets these were selected as developments sites rather than offsets. Arguably this concession places a barrier to achieving conservation for least cost, but if a planning system adheres to the irreplaceability metric, conservation outcomes are still safeguarded and eventually achieved. In total developers will bear a higher cost (**Fig.S12c**) than if conservation delivery was rigidly applied using SCP solutions, but the offset market allows both farmers and developers to choose where and when to participate.

In practice SCP solutions have rarely been implemented in their entirety because governments rarely have the authority or resources to implement a coordinated network of conservation sites simultaneously. Nevertheless, the principles of SCP such as complementarity underpin the irreplaceability metric, and why using it to guide offset markets strikes a balance between achieving conservation outcomes for least cost to society, while allowing developers the freedom to still select sites that offer the greatest potential profit to them. Likewise farmers individually retain the right to decide if they wish to participate. Implicit in a metric based on complementarity is a that offset site selection for conservation is as efficient as possible at the time a decision is taken, mean conflict with demands of developers and the farming sector are minimized as much as possible.

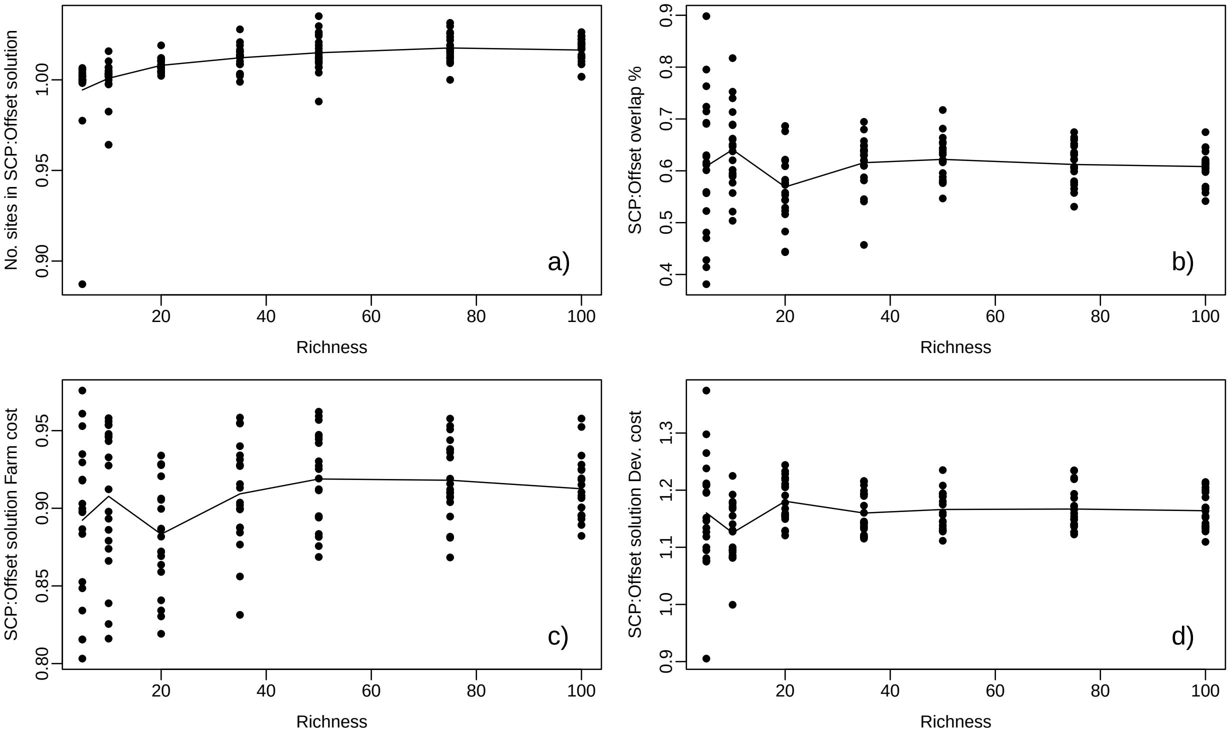

**Figure S12:** Relative efficiency of sites allocated to conservation by offset markets compared to systematic conservation planning (SCP) solutions in simulations with increasing species richness. Panel a) shows the ratio of planning units selected by SCP versus the offset-market, panel b) the proportion of planning units selected by the offset market that were shared with the SCP solution, c) the relative efficiency of the SCP solution compared to the offset-market based on the total cost of farmland conserved, and d) the total value of conserved sites for development using SCP solutions relative to the irreplaceability market cost-comparison.

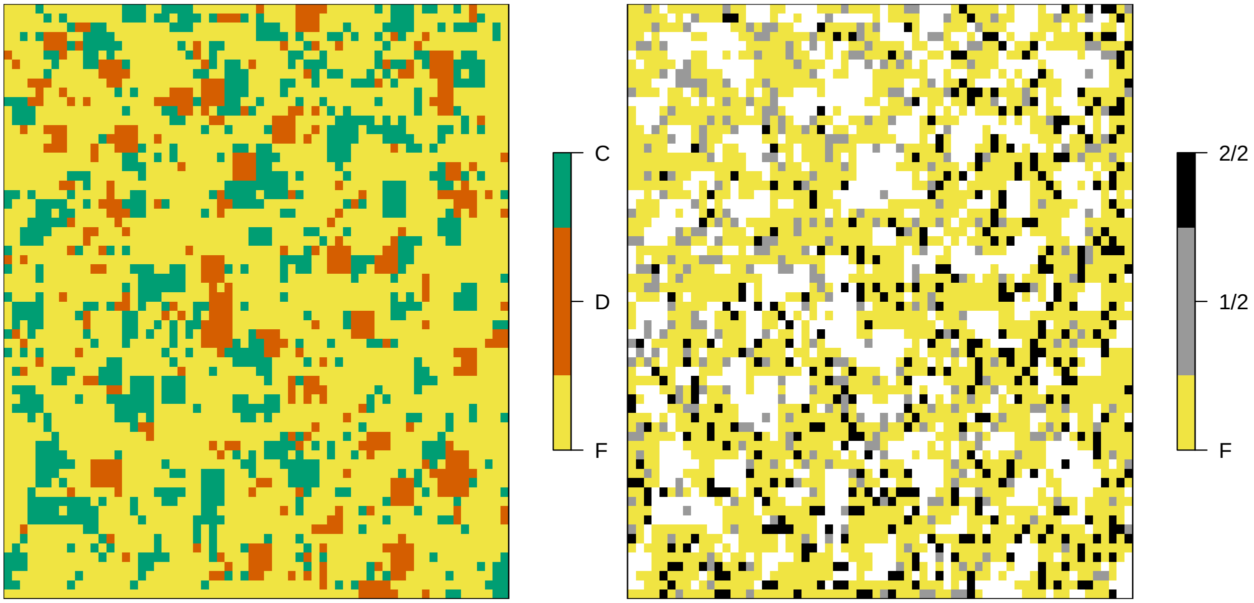

**Figure S13:** a) Initial state of the simulated landscape, divided into three classes: 70% farmland (**F**), 10% existing development (**D**), and 20% conserved (**C**). Map b) identifies the farmland selected for conservation by the optimal SCP solution and the sites selected by the offset market based on irreplaceability. Sites may be required by neither method, and remain farmland (**F**), or were selected by one (1/2), or both methods (2/2). The sites that began as existing development and conservation areas are excluded and appear white.

#### Comparison with Non-Systematic Offset Metrics

The comparison with systematic conservation planning above represents the *best-case* possible for achieving all conservation targets, but as discussed there are practical constraints that limit the implementation of those solutions in their entirety at once. The challenge that offset markets face when aiming to support conservation targets, which may equate to No-Net-Loss or Net-Gain, is therefore closer to other SCP tools that rank priorities to identify the *next best* action available for a given budget. However, a policy designer might prefer a simpler offset metric than a metric based on alpha-irreplaceability. To demonstrate that the benefits from using a metric founded on irreplaceability exceed those of an offset market based on other offset metrics we compared irreplaceability with three alternative site-based metrics that reduced the need for additional information.
> Offset Metric 1 (**OM1**) site scores were weighted by the inverse of each species range, and updated after each offset-trade, thereby favouring the rarest taxa in the landscape at the time.
> Offset Metric 2 (**OM2**) site scores were weighted by the inverse of each species range, but not updated.
> Offset Metric 3 (**OM3**) site scores were weighted by the number of species present.

OM1 assumes the same degree of knowledge as required for the summed alpha-irreplaceability, but without setting targets. Updates to planning units scores reflect species' global availability, but not complementarity to areas already protected. Offset metric 2 is equivalent to OM1, but values for each planning unit are not updated over time meaning weights for each species were fixed at their starting value. This metric required the same initial understanding of species distributions but does not require a register of species affected by previous offset transactions. Finally, planning unit value based on offset metric 3 (OM3) was based solely on how many species were present, but not as above, by which species, meaning only a map of species richness would be required to guide a market. OM3 thus represents the simplest offset metric considered here.

In general, non-systematic approaches to offset valuation were highly unlikely to achieve conservation targets before the market ceased trading (there were no longer any farmers Willing-to-Accept below the price at which developers were Willing-To-Pay). Based on a landscape with an even distribution of species, the desired targets were achieved by the offset market just once in 140 simulations using *Offset Metric 3*, 31 times using *Offset Metric 2*, and 3 times with *Offset Metric 1*. Most of the successful simulations were cases when the subset of species was 20 species or less. The success of these metrics in the landscapes that included moderate or steep gradients in species richness was even lower (24 and 5 of the 140 simulations respectively), and only for subsets of just 5 or 10 species. The failure of markets guided by the alternative Offset Metrics to achieve conservation targets is emphasized by **Figure.4** in the main text, and **Fig.S14** below. These figures show that in the majority of cases the markets exhausted the supply of farmers Willing-to-Accept, often allowing a far greater total number of developments. In landscape where richness becomes concentrated there is the opportunity for metrics like **OM3** to achieve target outcomes simply because features are co-located, but evidently this does not work well often enough, or for many species, and OM3 was least effective at achieving progress to species targets. Although neither **OM2** or **OM1** performed that well, especially for higher subsets of diversity, **OM2** actually performed better than **OM1**. This was because the targets we chose to demonstrate net gain with were all based on a very similar proportion of the starting potential, and because starting distributions of developments and conserved lands were approximately random, this meant that as **OM2** was guided by the starting rarity of each species this was a relatively effective proxy for the eventual targets. Normally conservation targets and the disparity between what was required relatively to what was available would be much more varied meaning this relationship would not hold as strongly. Although we might have assumed that updating the value of **OM3** would improve performance, doing so meant the market continually prioritized offsets at sites where the rarest species at the time were present. Subsequent trades would continue to prioritize the same species at the expense of achieving conservation outcomes for other species. This is why *complementarity*, understanding the marginal value of a new site against what has already been protected, is so important to systematic approaches.

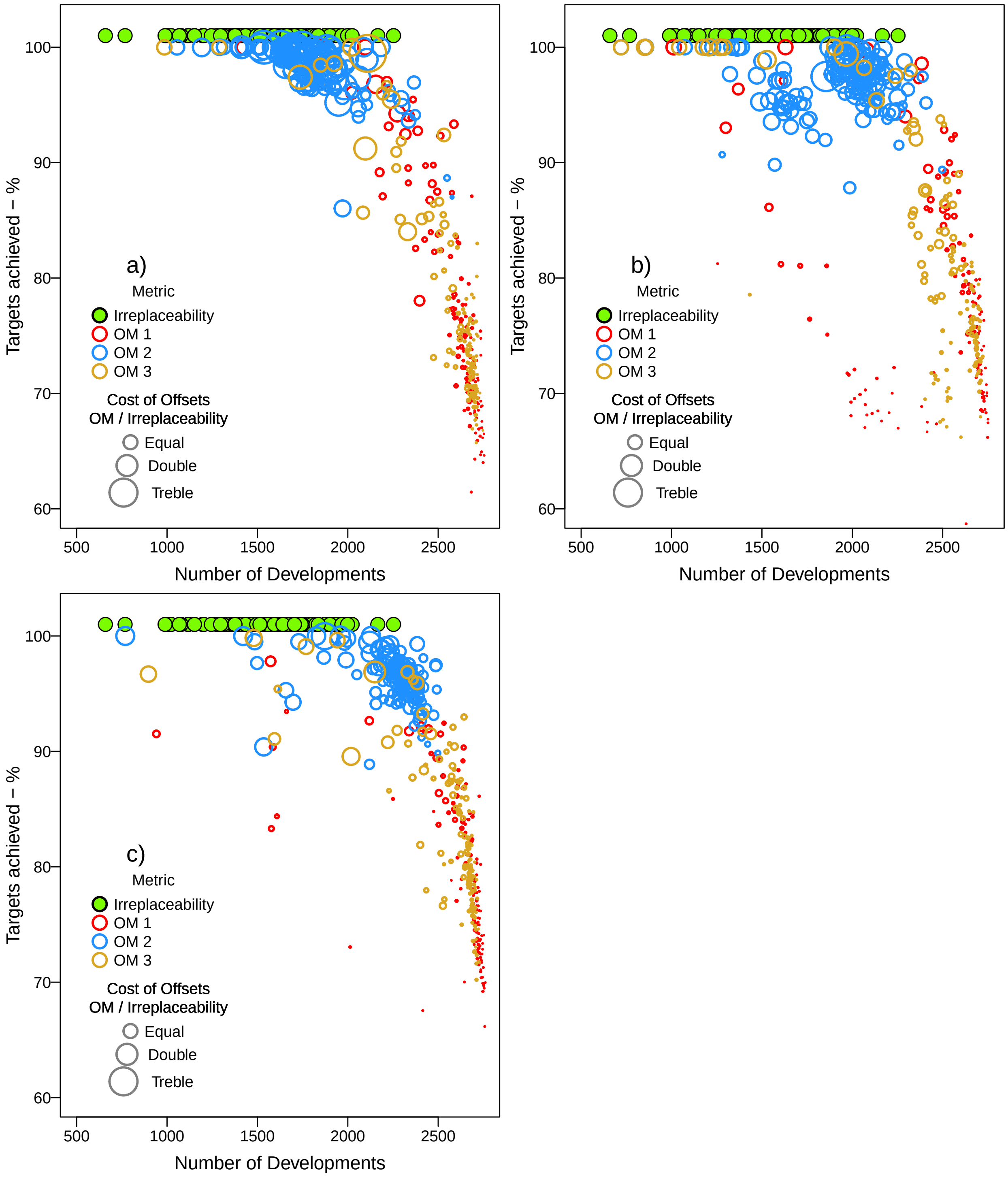

**Figure S14:** Performance of offset market simulations based on the proportion of conservation targets achieved and the number of developments required before market trading was ceased. Simulated markets guided by summed alpha-irreplaceability all ceased trading once all targets had been achieved (symbols offset to improve visibility). Simulations were guided by three further Offset Metrics, described in the main text (OM 1-3), and the size of those points further indicates the total cost of farmland required to offset developments relative to the cost of the equivalent solution using irreplaceability. Panels a, b and c provide the same analysis for the three landscapes with increasing bias in species richness.
